# Pipeline for retrieval of COVID-19 immune signatures

**DOI:** 10.1101/2021.12.29.474353

**Authors:** Adam J.H. Newton, David Chartash, Steven H. Kleinstein, Robert A. McDougal

## Abstract

**Objective:** The accelerating pace of biomedical publication has made retrieving papers and extracting specific comprehensive scientific information a key challenge. A timely example of such a challenge is to retrieve the subset of papers that report on immune signatures (coherent sets of biomarkers) to understand the immune response mechanisms which drive differential SARS-CoV-2 infection outcomes. A systematic and scalable approach is needed to identify and extract COVID-19 immune signatures in a structured and machine-readable format.

**Materials and Methods:** We used SPECTER embeddings with SVM classifiers to automatically identify papers containing immune signatures. A generic web platform was used to manually screen papers and allow anonymous submission.

**Results:** We demonstrate a classifier that retrieves papers with human COVID-19 immune signatures with a positive predictive value of 86%. Semi-automated queries to the corresponding authors of these publications requesting signature information achieved a 31% response rate. This demonstrates the efficacy of using a SVM classifier with document embeddings of the abstract and title, to retrieve papers with scientifically salient information, even when that information is rarely present in the abstract. Additionally, classification based on the embeddings identified the type of immune signature (e.g., gene expression vs. other types of profiling) with a positive predictive value of 74%.

**Conclusions:** Coupling a classifier based on document embeddings with direct author engagement offers a promising pathway to build a semistructured representation of scientifically relevant information. Through this approach, partially automated literature mining can help rapidly create semistructured knowledge repositories for automatic analysis of emerging health threats.

## INTRODUCTION

The rapid growth in scientific publications [1] presents a challenge for researchers to seeking a comprehensive understanding of the literature. This challenge is of particular importance in emerging disciplines and domains without existing comprehensive reviews or widely accepted frameworks for representing the field. The COVID-19 pandemic is one such example of an emerging publication phenomenon. While machine learning has provided many solutions for search problems related to information retrieval (IR) [2], application of IR to specific scientific domains remains an active area of research [3, 4]. Researchers have leveraged search engines to retrieve relevant literature, with keywords searches [5] or alerts [6], but these approaches usually require substantial further refinement.

Once relevant sources have been retrieved, information has to be obtained from the text. For some domains, machine consumable structures make specific data types trivial to extract, e.g. genes [7] and proteins [8], however integrating this information with a more comprehensive data model remains challenging. There are many methods to obtain salient information from identified sources, including; manual curation e.g. HIPC [5], rule-based semi-automated extraction of metadata from an abstract, e.g. the metadata suggestions for ModelDB [9], and PICO (population, intervention, control, and outcomes) extraction [10], which tags words related to the PICO elements in randomized control trials. Given the novelty of the scientific domain of COVID-19 research, it is difficult to known what information characterizes this subfield and how it will be presented in the paper. Thus, a semiautomated human in the loop approach facilitates a solution. COVID-19 may affect the human immune system in different ways. These effects – which could be at the level of changes of gene expression, of proteins, of metabolites, of antibodies, etc – may vary by population (e.g. young vs old), disease severity (e.g. mild vs severe), etc, with each pattern of effects constituting an *immune signature* for the disease. For some diseases (e.g. cervical cancer [11]), immune signatures have shown potential as predictors of survival or other clinical outcomes. Unfortunately, identifying papers containing human immune signatures and locating those immune signatures within publications is non-trivial. Immune signatures can appear in the text, figures or tables, with dozens of distinct signatures in a single publication, and may not be presented as the principle finding.

We developed a semi-automated pipeline (Figure 1), which utilized human-in-the-loop learning. As part of this pipeline we have created and validated a literature classifier that uses the abstracts and titles to retrieve papers likely to contain human COVID-19 immune signatures from a corpus of scientific literature. The pipeline then uses author solicitation: authors were asked to fill out a structured form describing the immune signature(s) in their papers. Author-supplied signatures from over thirty such papers are available on our website at covid-signatures.org.

**Fig. 1.**
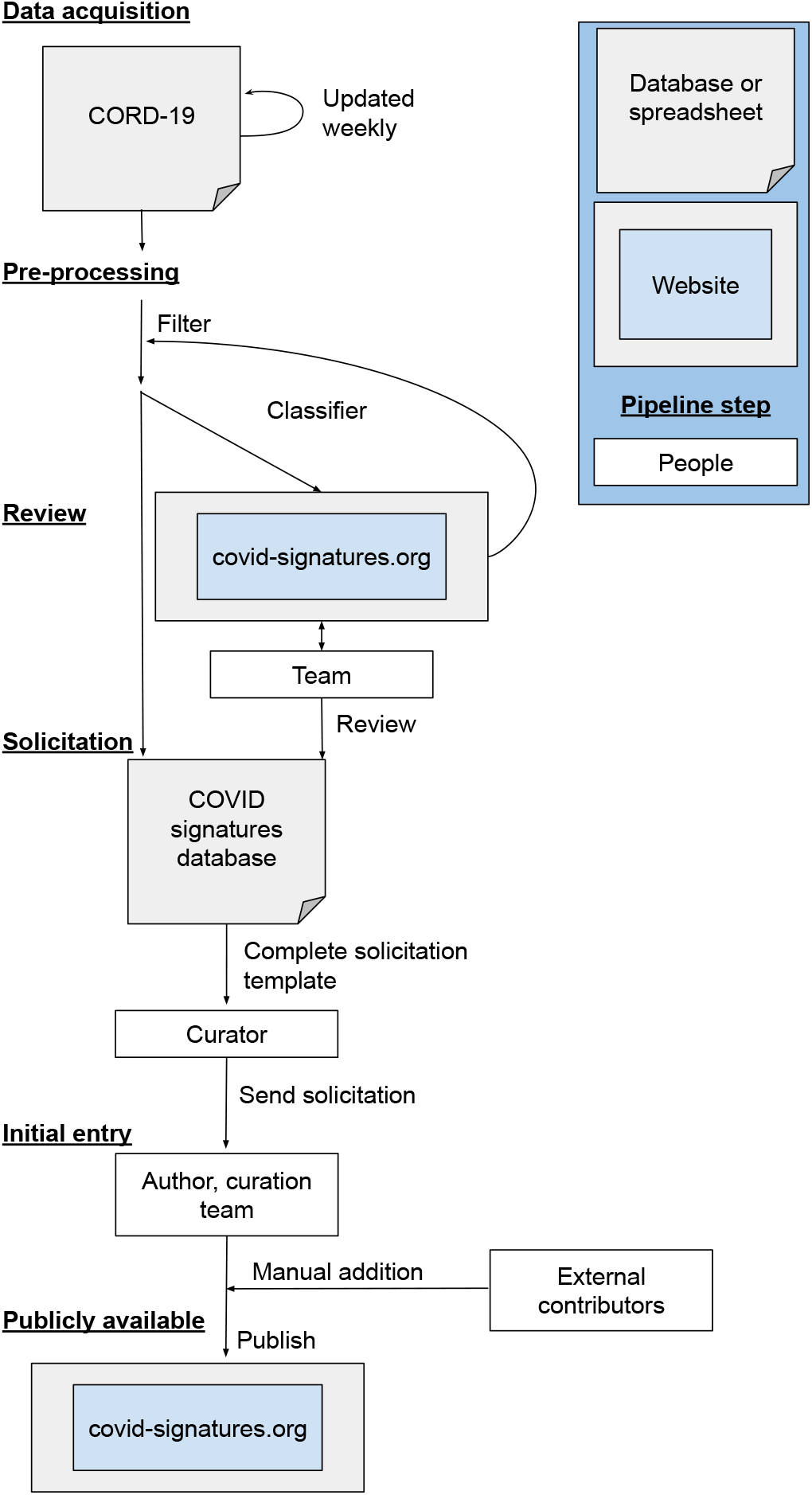
Pipeline for semi-automated curation of COVID-19 immune signatures. The steps in the pipeline correspond to those of a generic web service pipeline originally developed for ModelDB, a computational neuroscience model repository. Data acquisition utilized CORD-19 which required prepossessing; including substantially filtering of the dataset and removing duplicate entries. Triage divides the articles into one of 3 relevant or 2 not-relevant classes, which are then used to build a classifier to provide further articles for triage as well as directly for solicitation. Solicitation is done in a semi-automated fashion using an email template and an online form for the author to complete. Publicly available, after solicitation the information is made available via covid-signatures.org.

## MATERIALS AND METHODS

### Generic online platform

We developed a general purpose online literature review, author solicitation, and information sharing platform powered by the Django web framework (djangoproject.com) for templating and user management, a MongoDB database backend (mongodb.com), Bootstrap (getboostrap.com) for layout, and jQuery (jquery.com) for streamlined scripting (Figure 1). The pipeline logs timestamps and change history, including authenticated user IDs associated with the changes, to allow auditing and error recovery. To respect website visitor privacy, no cookies or other tracking mechanisms are sent to users that are not explicitly logged in; cookies are used to preserve authentication status across page loads. In particular, data providers who enter data using a special emailed link or via the unsolicited data entry process are not considered logged in and are not sent any browser tracking information. This generic platform is freely available at github.com/mcdougallab/pipeline. To adapt it for COVID-19 immune signatures, a JSON-encoded configuration file was used to specify database details, paper categories, explanatory text, data solicitation forms, email templates, etc.

### Reviewer interface

Expert review was performed using the aforementioned platform (Figure 1). A limited set of rules based on whole-word matching (e.g. the word “patients” implies that the paper studied humans) were used to tag the abstracts so that reviewers could examine by tag if desired (see supplementary material Table 1). Three expert reviewers each with at least five years graduate computational immunology training, examined the papers in the queue to determine whether or not they contained immune signatures and, if so, what type. The reviewers were presented with a title that links to the paper full text, abstract, selected metadata, and buttons to indicate their conclusions. To support reviewer corrections to automatic database population, an edit button allowed changes to the title and URL which were then pushed to the server via an AJAX call. For papers with a COVID-19 immune signature, reviewers were asked to choose from three broad classes of immune signatures. We included two additional review queues: “let’s discuss” for papers where the category was not obvious, and “review article” for work that may have a human immune signature but not be the primary source. An additional, auto-saved notes field allowed reviewers to make notes for themselves and for any future discussion. After 288 papers were reviewed, we tasked one of the expert reviewers with re-reviewing the papers to identify key words from the abstract that, in their judgment, made it more (e.g. “IL-8”) or less (e.g. “influenza”) likely that a paper would contain a COVID-19 immune signature. These identified terms were highlighted in the abstracts during the review phase (see supplementary material Table 2).

**Table 1.** Classifier confusion matrix based on LOOCV. Here the “present” class represents the presence of any type of human COVID-19 immune signature in the paper.

|  | predicted present | predicted not present |
| --- | --- | --- |
| present | 278 | 23 |
| not present | 49 | 142 |

**Table 2.** Second stage classifier confusion matrix based on LOOCV.. The correct predictions are highlighted in bold.

|  | Type A predicted | Type B predicted | Type C predicted |
| --- | --- | --- | --- |
| Type A signature | **42** | 15 | 1 |
| Type B signature | 6 | **135** | 18 |
| Type C signature | 2 | 45 | **27** |

### Data source

For interoperability with other COVID-19 literature analysis efforts through the use of shared identifiers, we leveraged the Allen Institute’s COVID-19 Open Research Dataset (CORD-19) [12]. CORD-19 provides a corpus with clear information retrieval benchmarks (see TREC-COVID challenge [4, 13, 14]), standardized machine readable data and SPECTER document embedding of each title and abstract [15]. To focus on primary sources that could contain COVID-19 immune signatures, we filtered this dataset to exclude:

- PubMed papers with “Comment”, “Review”, “Editorial”, or “News” article type.
- Papers from journals whose journal title includes “rev” as a whole word or as the start of a word to avoid review journals.
- Papers published before December 1, 2019.
- Papers that do not explicitly mention “COVID” or a related term (e.g. “2019-nCoV”) in the paper title or abstract.

CORD-19 is regularly updated (often weekly); we use these updates to add new papers to our pipeline. The results presented here were obtained with CORD-19 for the 8^*th*^ November 2021.

Duplicate and near duplicate papers frequently end up in the corpus, as many papers are released on preprint services before publication in a journal and CORD-19 include both preprint and journal publications. The CORD-19 Unique Identifier is linked to a conceptual document, which may include multiple versions of the manuscript. Near duplicate papers were identified using the SPECTER embedding with documents within a certain distance grouped together. Trial and error found 30 units in the SPECTER embedding space excluded most duplicates while preserving distinct articles. Within a given group, only the most recently released paper was used for further analysis. There are near duplicates in the corpus where one of the duplicates is missing metadata and the other is not, this can lead to one being filtered from the dataset by our preprocessing, while the other is not. For this reason, we also remove entries with near duplicates in our excluded set.

### Two-stage SVM classifier

We developed a two-stage Support Vector Machine (SVM) based classifier for the filtered CORD-19 literature, using `sklearn`version 0.24.2 [16] with Python 3.6.10. For the first stage, the SVM model (polynomial kernel of degree 4) simply seeks to determine if a paper contains an immune signature or not; this SVM was trained by grouping the three immune signature classes into one super class and the “review article” and “no signature” classes into another super class. A second SVM model (polynomial kernel degree 5), trained on only the papers confirmed by our expert reviewers to contain a COVID-19 immune signature, was used to predict the type of immune signature that would be present for those papers predicted by the first classifier to contain an immune signature. Probabilities were obtained from the SVM classifiers using Platt scaling [17]. Both SVM models were trained and applied on the SPECTER embeddings of the title and abstract, not directly on the text. We store both the paper metadata and the predicted probabilities in the database.

### Immune signature contribution from paper authors

Once papers containing COVID-19 immune signatures have been identified, either manually or automatically, we contacted the corresponding author(s) to request details of the immune signatures in the paper, corresponding to our data model Figure 2. Our platform provides a form with a unique URL for each entry in the database. This form identifies two classes of data – one that is global and applies to the entire form – and one that pertains to a specific fact about the paper (in our case, a specific immune signature), of which there can be many. The field names are configurable via the JSON configuration file, but for this project, the form asks for global data identifying the paper and contributor (reference, contributor, organization, and email address), and specific instance data about each immune signature (description, location in the paper, tissue, immune exposure, cohort, comparison, any repository ID, analysis platform, response components and response direction).

**Fig. 2.**
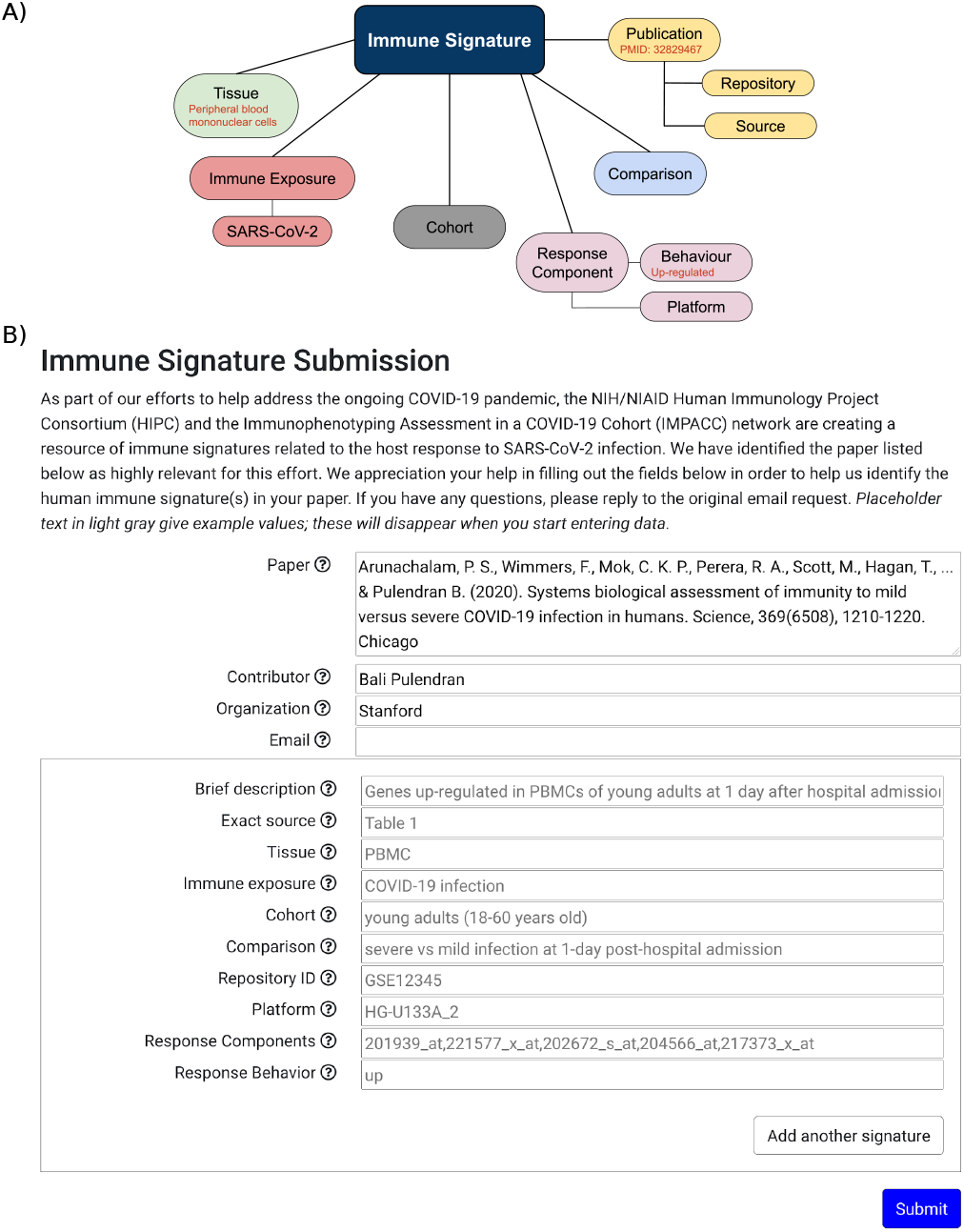
Immune signatures model and solicitation. **A)** Our data model for COVID-19 immune signatures is based on [5]. **B)** The immune signature template provided to authors, with help text shown in gray.

Once the global data is entered, a button on the data entry form (visible to authorized, authenticated users only) generates an email from a template and the field data. The email recipients receive a link to a page for just their specific paper, which does not require a login. Each field on the form has associated help text and examples. All fields are editable by the email recipient except for the paper reference. An arbitrary number of immune signatures may be entered for each paper. When the contributors press the “submit” button, the entered data is stored in the database and logged in a separate file, allowing administrators to revert to a previous version in the case of accidental or malicious changes after initial data entry. A typical data entry form is shown in Figure 2.

To allow third-party manual solicitations, a “submit your immune signatures” button on the covid-signatures.org homepage opens an entry form that is the same as one seen by solicited contributors except without the pre-filled global fields and with an editable paper reference field. These entries are assigned an automatically generated internal identifier which the website administrators can later map to a CORD-19 identifier.

### Data dissemination

The contributor-entered details of the immune signatures are stored in a MongoDB database and are made available via covid-signatures.org in both HTML and JSON. Internal users can access the full database entries, whereas the public version does not include edit history, contributor details, or internal identifiers (but does include the CORD-19 identifiers).

### Data Analysis

Python Data Analysis Library `pandas`0.24.2 [18] was used to manage the data from CORD-19 and the pipeline for analysis. We used `sklearn`0.24.2 [16] to perform k-means clustering, for SVM and logistic regression classifiers with tolerance set to 10*−*7. Uniform manifold approximation and projection (`UMAP`) 0.4.6 [19] was used to visualize the clusters. `SciPy`1.5.4[20] was used for statistical tests. To evaluate the classifier, we used Natural Language Toolkit (`NLTK`) 3.5 [21] for tokenization, excluding English stop words and words with less than three characters. WordNet [22] was used to lemmatize the words. TD-IDF computation was facilitated by the `sklearn`package, including 1-and 2-grams. The logistic regression classification used inverse of regularization strength 50. When comparing word frequencies, we excluded words that occurred fewer than five times in the titles and abstracts of the selected papers. We used `Gensim`3.8.3 [23] to perform Latent Dirichlet Allocation (LDA) [24] with 1000 iterations and 100 passes, on the filtered CORD-19 abstracts and titles, excluding words that occurred in more than 80% of abstracts or fewer than 5%.

## RESULTS

Our overall workflow involved the development of a training set of papers containing COVID-19 immune signatures, SPECTER and SVM-powered identification of papers likely to have these immune signatures, expert review of a subset of papers, data solicitation from the authors, and then data dissemination on the covid-signatures.org site (summarized in Figure 1). As we envision this workflow will generalize to other semi-structured data acquisition efforts, we developed a generic online platform to streamline its application.

### Papers with immune signatures are clustered in SPECTER abstract embedding

We sought to determine whether SPECTER embedding preserved sufficient information to identify papers containing COVID-19 immune signatures. SPECTER provides document-level embeddings using a pre-trained language model. The CORD-19 dataset provides a SPECTER embedding, a 768 dimensional vector representation of the title and abstract, for each of the papers. We manually identified 5 papers from CORD-19 with COVID-19 immune signatures. We also considered an additional 69 papers with non-COVID immune signatures that were identified as part of the Human Immunology Project Consortium (HIPC) [5], together with a matched control group of papers for the HIPC immune signatures taken from the same volume of the journals. We used the pre-trained SPECTER model[15] to obtain an embedding for each of the additional papers from their title and abstract. *K*-means clustering applied to the SPECTER embeddings identified *k* = 6 clusters based on Akaike Information Criterion (AIC).[25] Almost all of the papers with immune signatures were grouped in a single cluster, cluster 3. Four of the 5 papers with COVID-19 signatures and 68 of 69 papers with vaccination signatures were in cluster 3. The remaining papers with immune signatures were part of a second cluster, cluster 6 (1 of 5 papers with COVID-19 signatures, and 1 of 69 papers with vaccination signatures). In contrast, a control group of papers were more widely dispersed in the embedding space with a significantly different distribution compared to the papers with vaccination signatures (*χ*^2^ = 14.42, *p* = 0.006) (Figure 3).

**Fig. 3.**
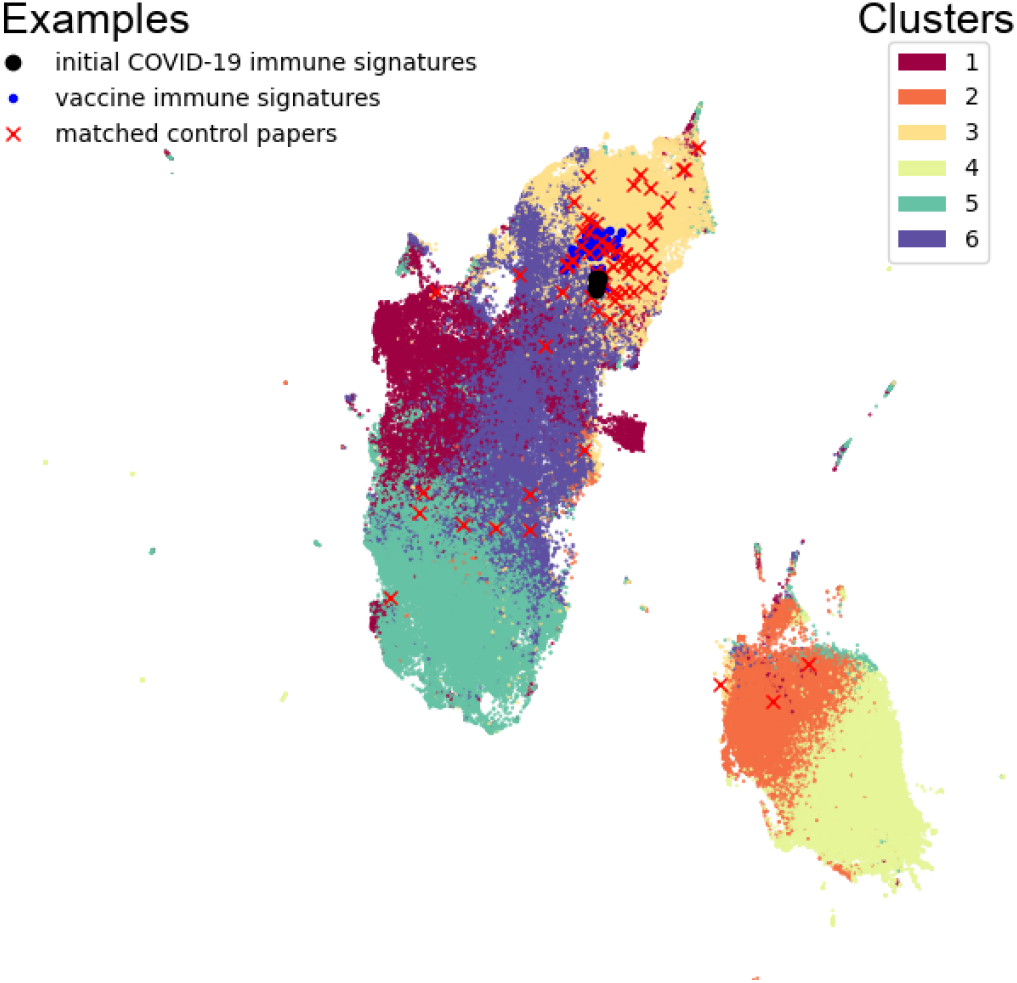
Uniform Manifold Approximation and Projection (UMAP) of *∼* 170, 000 papers from the CORD-19 dataset (small points) based on their SPECTER embedding. Manually selected papers with COVID-19 immune signatures (*n* = 5; large points), vaccine response immune signatures (*n* = 69; * markers), and matched control papers (*n* = 68, x markers) are marked. Each paper was assigned to a cluster via *k*-means clustering, with *k* = 6 based on the AIC.

This significant sub-clustering suggests that the SPECTER embedding effectively preserves information on whether or not a paper contains an immune signature. These results suggest that we can use the SPECTER embedding as the basis to construct a classifier to identify papers with COVID-19 immune signatures.

### Papers with COVID-19 signatures can be predicted with high accuracy

We built a classifier to determine if the SPECTER embedding could be used to predict which papers contained COVID-19 immune signatures. To create the classifier we bootstrapped a training set, consisting of papers with a label indicating whether they contained immune signatures. We took advantage of the close grouping of immune signature papers in the SPECTER embedding by starting with five initial papers with COVID-19 signatures (shown in Figure 3), and identifying the nearest 100 papers to each of these points in the embedding space. We also included the 100 papers from the filtered CORD-19 dataset that were nearest to the center of the vaccine immune signatures.

These papers were labeled by expert reviewers, using the pipeline interface. To allow simultaneous article review by multiple parties, only a random, small portion of the dataset is shown on the pipeline review platform at a time by default, minimizing the risk of duplicate review of the same paper. Overall, this identification and review process resulted in 271 papers, of which 140 contained immune signatures. An additional 52 papers from CORD-19 identified by rule-based filters were subjected to expert review, identifying another 11 relevant papers.

### Inter-rater reliability of signature presence

To test the reliability of our expert reviews, we selected 100 articles at random from the set of papers to be reviewed independently by two reviewers. The two reviewers had 92% agreement on determining whether a paper contained a COVID-19 immune signature. Cohen’s kappa coefficient [26], a robust measure of inter-rater reliability that accounts for chance agreements, shows very good agreement (0.84, 0.73-0.95 95% confidence interval). Papers where reviewers did not agree on the presence of an immune signature were not included in the training set for the classifier.

### Classifier bootstrap and performance

A SVM classifier [16] was fit to the 316 papers unambiguously identified by reviewers as either containing or not containing immune signatures. To achieve reliable classifier predictions, we iteratively selected additional papers for review based on the prediction of the classifier. We added papers with estimated probability of containing human COVID-19 immune signatures between 0.80 and 1.0 to increase the number of positive examples (Figure 4). This iteration was deemed sufficient as the marginal improvements to the classifiers performance were small. Specifically, the average reduction in positive predictive value (PPV) based on Leave-One-Out Cross Validations (LOOCV) with one fewer paper was 0.10 *±*0.28 (from 81.3% to 81.2%). Overall, this yielded a set of 216 papers containing immune signatures and 185 papers without immune signatures.

**Fig. 4.**
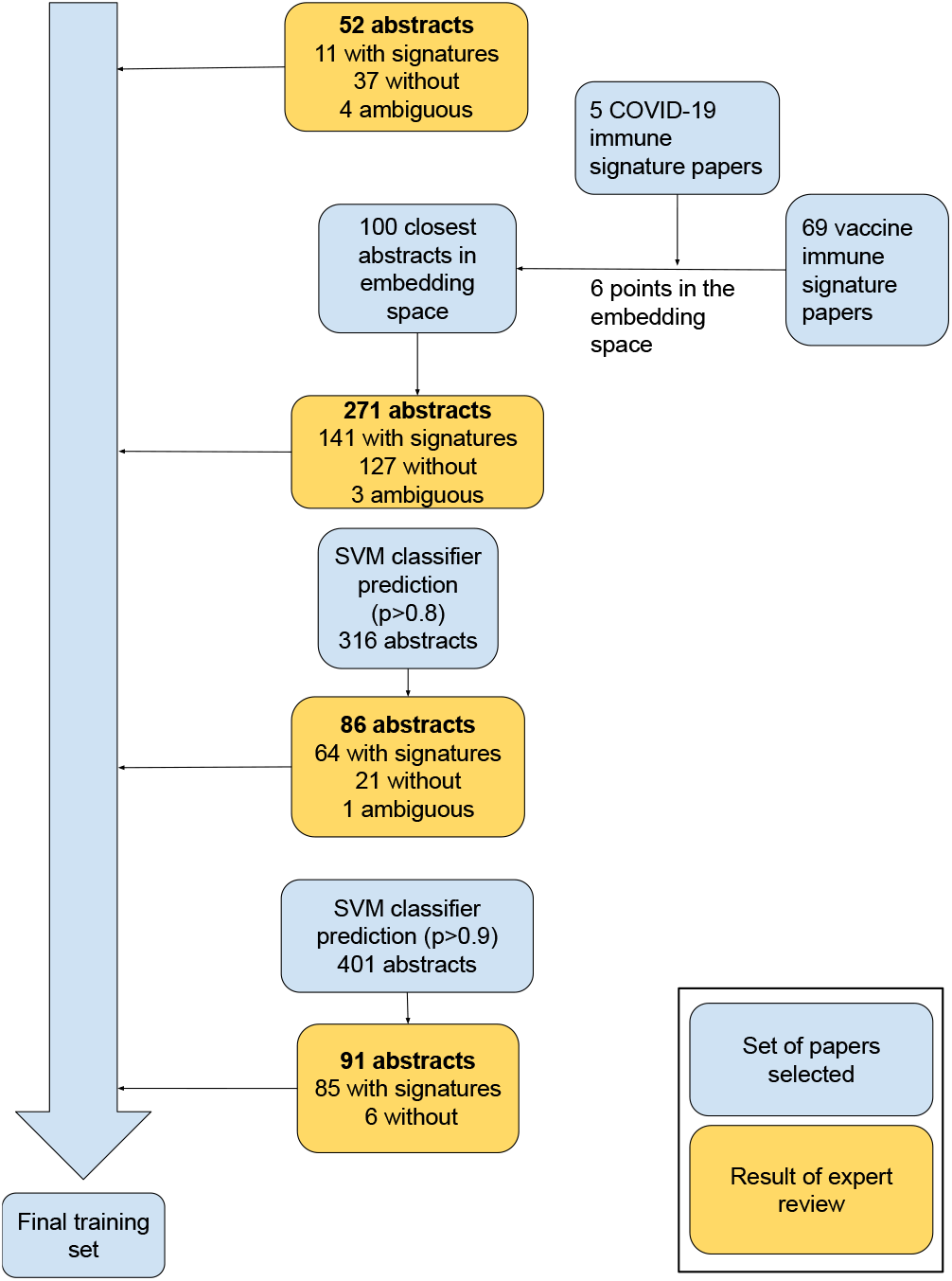
Iterative identification of papers for bootstrapping the classifier. The bootstrapping process was initiated papers selected with rule-based filters for expert review. We then leveraged the SPECTER embedding to select papers for review based on their distances in the embedding space. Once an initial training set was bootstrapped, we fitted an SVM classifier with the 316 abstracts and use it to select 86 additional papers. An SVM classifier fitted to the 401 abstract, predicted 91 highly likely papers to validate the classifier. This gave a final training set of 492 papers, 301 with immune signatures, 8 of which were ambiguous (where reviewers did not agree on the presence or absence of immune signatures).

Using a SVM classifier trained on these 401 papers, we selected an additional 91 papers based on predicted probability (*>*= 0.8) to be highly likely to contain a COVID-19 immune signature. An expert reviewer (SK) then classified these articles, and determined that the majority (92%) contained immune signatures. Given the size of the corpus, and the fact that emailing corresponding authors included a manual step (i.e., a human-in-the-loop, see methods), we decided to use a greater specificity threshold to select papers for direct solicitation in order to increase efficiency. Considering a higher threshold (predicted probability *>*= 0.9), of the 61 papers that met this threshold, 60 of them contained COVID-19 immune signatures (94%). These additional papers gave a final training set of 492 papers, with 301 containing COVID-19 immune signatures.

We evaluated the performance of the SVM classifier on our training set using LOOCV. The receiver operating characteristic (ROC) area under the curve (AUC) was 0.916 (Figure 5A). With a probability threshold of 0.5, the SVM classifier had a PPV of 86%, with accuracy 85%, sensitivity 92%, selectivity 74% and F1-Score 89% (Table 1). Applying this SVM classifier to the filtered CORD-19 collection identified (at the probability threshold of 0.5) around 15,500 papers (*∼* 2% of the CORD-19 corpus) are likely to contain COVID-19 immune signatures.

**Fig. 5.**
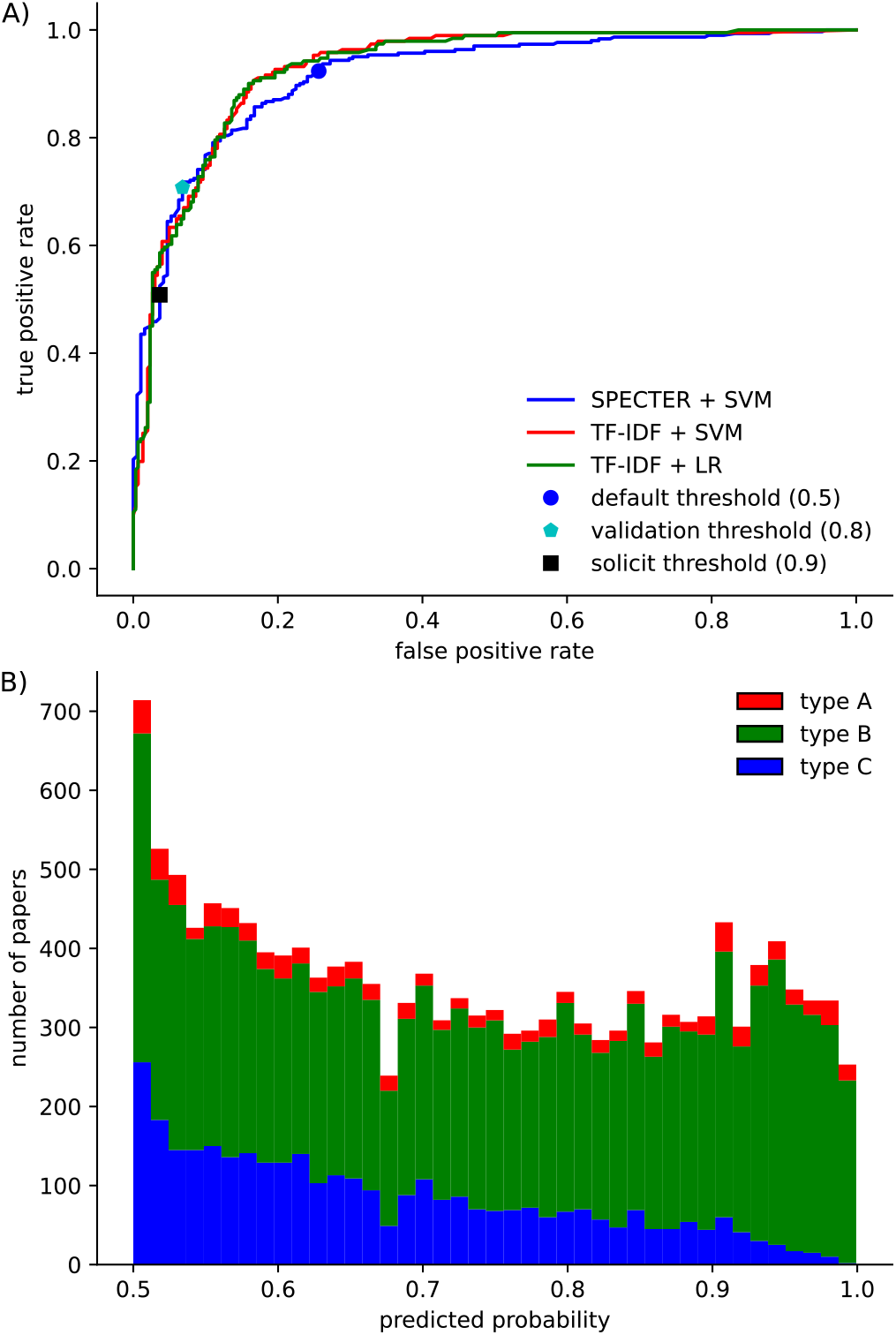
The classifier can identify papers containing immune signatures from the SPECTER embedding. **A)** The receiver operating characteristic (ROC) curve based on LOOCV for the classifier with the default threshold (0.5), validation threshold (0.8) and direct solicitation threshold (0.9) highlighted. Different lines are shown for the SVM classifier with SPECTER, SVM classifier with TF-IDF and LR with TF-IDF. **B)** Number of papers in the CORD-19 dataset predicted to have immune signatures of different types at different thresholds of the classifier. The first SVM classifier predicted the probability of the paper containing an immune signature at the threshold indicated. The second SVM classifier then predicted the type of immune signature.

We evaluated whether the SPECTER embedding was essential for achieving the high performance of the classifier. We compared the SVM classifier using the SPECTER embeddings to the widely used approach of Term Frequency Inverse Document Frequency (TF-IDF) [27] of the titles and abstracts, with both a SVM classifier and with logistic regression. TF-IDF with SVM classifier achieved an AUC of 0.928 using LOOCV, which was very similar to the SPECTER approach. Likewise TF-IDF with logistic regression had similar performance with an AUC 0.928 (Figure 5A). As these alternative approaches performed similarly, we chose to use the SPECTER embedding for the rest of this study as it was supplied with the CORD-19 dataset and did not require any additional text processing.

### The SPECTER embedding captures information about the type of COVID-19 signature

We next sought to determine if the SPECTER embedding of titles and abstracts contained sufficient information to distinguish between papers describing different types of COVID-19 immune signatures. We divided the COVID-19 immune signatures into three types: (A) Type A papers contained gene expression signatures, (B) Type B papers included signatures involving proteins, metabolites and/or cell types, and (C) Type C papers included all other COVID-19 immune signatures.

### Inter-rater reliability of signature type

To verify that the different signature types were meaningful and distinct, two experts reviewed a set of 100 papers. Papers where reviewers agreed on the type of immune signature (38 of 46) were added to the training set for the second stage classifier. The classes of immune signatures were well recognized by our reviewers, with substantial (82%) agreement when including whether a paper contained an immune signature or not and the type of immune signature (*κ* 0.72 0.61-0.83 95% confidence interval) (Table 3).

**Table 3.**
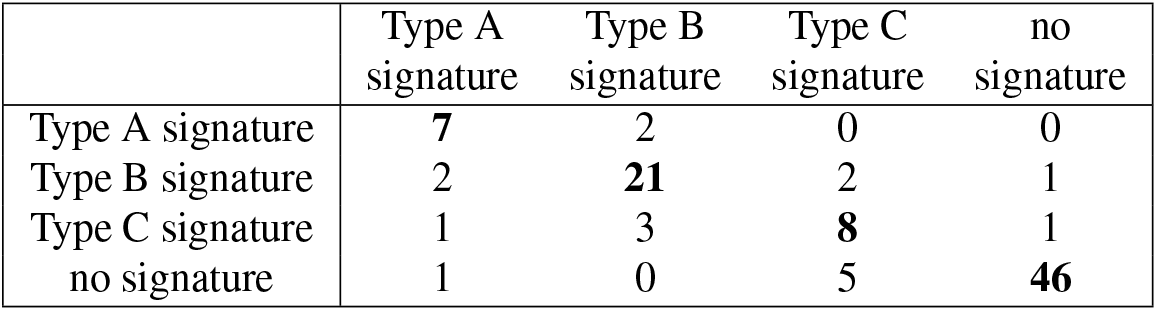
Independent expert review of articles showed substantial agreement between the three immune signature classes.

|  | Type A signature | Type B signature | Type C signature | no signature |
| --- | --- | --- | --- | --- |
| Type A signature | **7** | 2 | 0 | 0 |
| Type B signature | 2 | **21** | 2 | 1 |
| Type C signature | 1 | 3 | **8** | 1 |
| no signature | 1 | 0 | 5 | **46** |

### Classifier performance

Using the papers that were initially identified as containing COVID-19 immune signatures, we fit a “second stage” SVM classifier to predict the type of signature. To evaluate performance, we tested the classifier using the 84 of the 91 papers used to validated the previous classifier. The 91 papers were predicted to contain immune signatures (probability threshold *>*= 0.8) and the 84 used here are those that were validated by expert review as actually containing immune signatures. These papers included all three of the signature types: 10 type A, 58 type B, and 16 type C signatures. On this independent test set, the weighted averages (where each class’s contribution is scaled by the fraction of papers in that class) were PPV 74%, with sensitivity of 58%, specificity 58% and F_1_ score 61%. Incorporating these newly reviewed papers provided 291 annotated papers with immune signatures (out of the 506 total papers in our final training set). We then refit the classifier to the 291 papers with signature types. LOOCV of this “second stage” classifier (Table 2), gave weighted average PPV 69% with sensitivity 70%, selectivity 80%, and F_1_-Score 69%. The classifier performed best on “Type A” signatures, while the broadest category “Type C” had the most incorrect predictions. These performances strongly suggests that the embedding is able to capture the the differences between signature types.

### Features driving the classification

To discover which features our classifiers were using to determine the presence of immune signature, we compared papers with high and low probabilities of containing COVID-19 immune signatures. For the high probability papers, we selected the 500 articles with the highest predicted probability to contain an immune signature based on the classifier. As a comparison group, we selected 500 papers with a low probability (*∼* 0.1) of containing an immune signature. This low probability was chosen for the comparison set to avoid selecting marginal papers from the corpus and those not written in English. A log-likelihood comparison of word frequencies [28] between these two groups identified 171 words with significantly different frequencies, of which 38% were more frequent in papers containing immune signatures (*χ*^2^ with adjusted 5% threshold for 876 comparisons). The top 10 differences are shown (supplementary material Table 3) and are consistent with the focus of the classifier on human immunology, e.g. “patient”, “health”, “cell” and “severe”.

To further interrogate the classifier, we applied LDA topic modeling to the CORD-19 corpus filtered to include only potential relevant entries as described in methods (Figure 6). When comparing the sets of papers described above, papers predicted to contain immune signatures were predominantly related to topics 6 and 8, which appear to describe immunological and clinical work. There were significant differences in the topic composition across all topics (Mann-Whitney U tests at the Dunn–Šidák corrected 5% significance level for 10 tests).

**Fig. 6.**
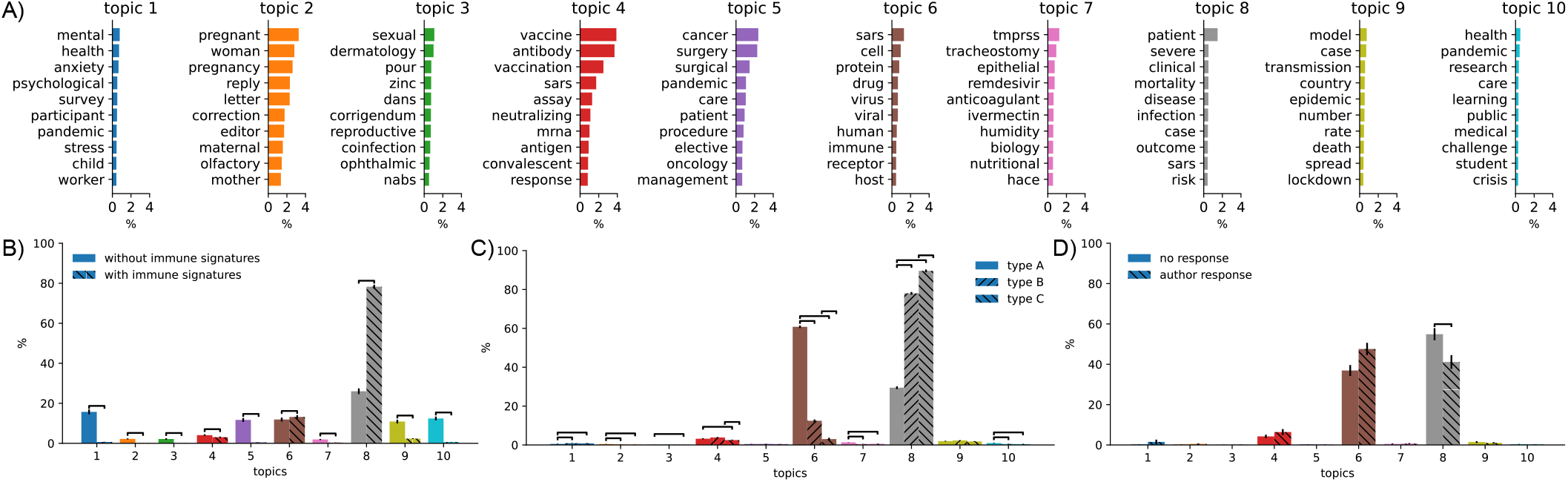
LDA of the CORD-19 dataset with average scores given to a samples of papers based on SVM classifier predictions. **A)** 10 topics identified from the abstracts and titles of the filtered CORD-19 corpus. **B)** comparison of the average topic composition for a sample of 500 papers predicted to contain COVID-19 immune signatures and 500 papers predicted not to contain immune signatures. **C)** comparison of average topic composition for three samples of 500 papers predicted to contain a specific type of COVID-19 immune signature. **D)** comparison of average topic composition for the 31 solicited papers where authors provided immune signatures and the 69 where they did not. Error bars show the standard error of the mean. Statistically significant differences between topics is shown based on Mann-Whitney U tests at the 5% significance level using the Dunn–Šidák correction for multiple tests.

Next we consider which features may be driving the “second stage” classifier in assigning different signature types. We chose a random sample of 500 papers most likely to contain a particular immune signature type (prediction *p >* 0.5 of containing immune signatures and *p >* 0.5 for the type of immune signature). Comparing word frequencies between the titles and abstracts of the different signature types identified 413 statistically significantly different words for type A COVID-19 immune signatures (26% had higher relative frequency in type A sample). Similarly for the predicted type B titles and abstracts there where 343 statistically significantly different words (14% with greater relative frequency in the type B sample) and 326 for type C (with 15% with greater relative frequency in the type C sample). For each case, the ten words that differed the most are shown (see supplementary material Table 3), with bold typeface indicating they occurred with greater relative frequency in that sample. These word frequencies suggest one way that the classifier predicts the paper contains an immune signature may be the presence of certain words associated with immune signatures (e.g. “patient”, “cell”, etc). In contrast, the type of immune signature may be determined by a reduction in the frequency of confounding words, such as “patient”, “clinical”, etc for type A immune signatures. We performed LDA topic modeling for different types of immune signatures. While papers with all types of signatures were predominately associated with topics 6 and 8, there were significant differences between them, with type A being 61% topic 6, type B 12% and type *C* only 3%.

### High author response rate was found for immune signature extraction

Previous work demonstrated many authors are willing to provide both data and metadata about their work [9]. To discover if this is also the case for COVID-19 immune signatures, we developed a semi-automated pipeline to contact the corresponding author(s) of papers and ask them to provide details of their published immune signatures via an online form (Figure 2).

Articles were chosen via expert review or based on a predicted probability from the SVM classifier. Expert review is the gold standard for the presence of immune signatures, when using the classifier, to avoid an unnecessary burden corresponding with authors of potentially unrelated work, we required a higher specificity when retrieving papers for solicitation. We used probabilities *>* 0.9 for containing an immune signature (from the first SVM classifier) and combined probability *>* 0.8 of it being type A or B (from the second classifier). The high threshold for the predicted probability of having any immune signature changes the confusion matrix and subsequently increases the specificity to 96% at the cost of sensitivity (50%), with PPV 96%. Although the SVM classifiers were used to select articles for direct solicitation, a human was kept in-the-loop for quality control and validation. We selected a convenience sample of 100 papers to directly solicit information on the COVID-19 immune signatures (63 of these papers were selected by the classifier and 37 by expert review).

We sent solicitations to authors of the 100 selected papers; 31% of these authors contributed immune signature information. Notably, we sent only a single solicitation email to each author with no reminders. The majority of responses (20 of 31 papers) submitted a single immune signature, while 2 signatures were specified in 5 of the responses, 3 signatures were specified in 2 responses, and one contributor each specified 4, 12, 14, and 27 signatures for a single paper. While the submission form did not have any required fields, all of the responses included a free-text description of the signature, the comparison underlying the signature, and the source of the signature in the paper (e.g., the figure or table number). However, only 35% (11 or 31 responses) provided a list of the immune response components (e.g., gene or protein names) that comprise the signature. Thus, while the overall response rate to direct solicitation was reasonable, follow-up is needed to obtain signature details.

We next sought to determine if there were differences between the papers the led to successful solicitations compared with those that did not. The likelihood of obtaining an authors response for the 63 papers selected with our classifiers was not correlated with the predicted probability of the paper containing an immune signature (Point-biserial correlation coefficient coefficient *r* = *−*8.09 *×* 10*−*4, *t*_61_ = 6.32 *×* 10*−*3 *p* = 0.995). The lack of correlation may be because of the small probability range chosen for solicitation. We did not detect a difference in the predicted class (“Type A”, “Type B” or “Type C”) for papers with or without author responses 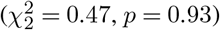. Finally, although the response rate of papers chosen by the classifier was lower than those chosen by expert review (27% vs. 38%, respectively), this difference was not statistically significant 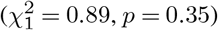.

Topic modeling of the papers where we solicited further information showed they were predominately topics 6 and 8 (Figure 6D). There were significant differences between the contribution of topic 8 where authors responded compared with papers where authors did not, 5% level with Dunn–Šidák correction for 10 comparisons (Topic 6; MannWhitney U p=0.015, Topic 8; p=0.0045).

## DISCUSSION

We used SPECTER and machine learning to analyze titles and abstracts of papers from CORD-19 to predict the presence or absence of COVID-19 immune signatures and the type of immune signatures. For papers predicted to contain immune signatures, we solicited additional details from the corresponding authors, and disseminated the information received on the covid-signatures.org website.

The method presented here is an adaption of our generic pipeline to focus on obtaining COVID-19 immune signatures Figure 1. This generic approach was originally developed in our group to help populate a computational neuroscience model repository, ModelDB [29]. As with COVID-19 immune signatures, model data from computational neuroscience papers cannot be obtained from the abstract, despite presence being retrievable. We believe that this pipeline approach would work for other categories of research. The willingness of authors to share information, however, likely varies by field and over time as the culture of the field shifts (see e.g. the discussion in [30]).

By providing a similarity metric, SPECTER allowed us to build a corpus starting from only five examples of papers with COVID-19 immune signatures. The utility of the SPECTER model is due to its general purpose representation of the scientific literature, trained on the Semantic Scholar Corpus [31] of over a hundred thousand articles with their citation information. TF-IDF with logistic regression was comparable to the generic SPECTER given sufficient training data. The SPECTER model could be further trained to specialize it to a given corpus, such as the COVID-19 literature, possibly improving the embedding and classifier performance in this field.

To evaluate which features might be driving the classifier we consider the word frequencies and performed LDA topic modeling (Figure 6). Word frequencies suggested the type of immune signature may be determined by the absence of confounding words, particularly for the broader signature types B and C, where the the significant differences in word frequencies were mostly due to words occurring less frequently (86% of the significant different words for type B and 85% for type C). LDA topic modeling showed papers with immune signatures were mostly represented by the clinical and immunological topics and had a significantly different distribution of topics compared to those without immune signatures. There were also significant difference between the topic distribution of the types of immune signature. Type A signatures were primarily represented by the immunological topic followed by the clinical topic, this relationship was reversed for type B and C signatures.

Author solicitation is an important part of the pipeline, and we found many (31%) authors were willing to provide additional details about their research. This high response rate was achieved despite there being no public platform to demonstrate how it would be used during the development of this work. Both the author response rate and the quality of authors responses are important for the utility of our pipeline approach. To reduce author burden and enable direct linkage of the signature to text in the manuscript, we requested authors report information as it appeared in the paper rather than attempting to translate terms to a standardized form (e.g., leveraging the Cell Ontology for cell types that are reported as response components) [32]. Thus, signatures included references to “NK cell”, “CD3 T cells” and “gammadelta T cells,” which could then be mapped to terms from the Cell Ontology: “natural killer cell” (CL:0000623), “T cell” (CL:0000084), and “gamma-delta T cell” (CL:0000798), respectively. This gives us a data set that represents immune signatures as they are likely to appear in publications, which can assist in future development of methods to detect immune signatures and their component parts.

We focused the machine learning and natural language processing on the paper titles and abstracts, but there is more information available in the full text and, in particular, the COVID-19 immune signatures are themselves expressed in the body of the paper (figure captions, results, etc). However, the structured full-text is only available for about a quarter of the CORD-19 dataset and there is lower signal-to-noise ratio in the full-texts than in abstracts [9, 33]. Nonetheless, some types of data are relatively unambiguously identifiable from the full text. In particular, journals often require certain types of data to be archived in a repository, such as GEO, ArrayExpress and FlowRepository. These repositories use standardized formats for their accession IDs that can be identified using regular expressions. However if the paper contains multiple signatures in multiple repositories, manual curation is still necessary to correctly divide them between immune signatures.

More sophisticated language models facilitate higher performance on information extraction tasks, e.g. SciBERT (which underlies SPECTER) has shown success in retrieving Population Intervention Comparator Outcome (PICO) elements in the randomized controlled trial literature [10]. The PICO elements have been tested to both identify text spans and extract structured information for automated biomedical evidence extraction. [34] While PICO elements partially overlap the immune signature data model (Figure 2), there are several challenges to leverage PICO extraction to further automate COVID-19 signature extraction. For example, information about immune signatures is often not present in the text, but rather in figures or tables [5]. Current PICO extraction methods also group the intervention and comparison categories, which makes it more difficult to apply them to our data model. Papers containing multiple signatures present the additional challenge of grouping the PICO elements for a particular immune signature. PICO predictions could be used to assist manual entry, such as by providing paper-specific autocomplete suggestions or placeholder text to reduce the burden of data entry. This approach risks, however, biasing the author responses. The author-supplied information may be used to improve machine learning for identifying PICO information by providing data for model training and testing, although this may be complicated by the variability with how author contributors specify their COVID-19 immune signatures.

Crowd-sourcing further curation offers a potential mechanism for reducing variability and providing consistently annotated human COVID-19 immune signatures similar to those made available through the HIPC Dashboard [5]. A portal is under development that facilities community annotation of COVID-19 immune signatures community.covidsignatures.org/, utilizing the signatures retrieved by this pipeline.

Identifying papers containing COVID-19 immune signatures and collecting data and contextual information (metadata) regarding the signatures can help speed scientific progress in this area. As has been seen with vaccination [35] and inflammation [36], providing access to such a resource supports secondary and comparative analyses resulting in a broader understanding of immune system response. We have demonstrated this pipeline approach Figure 1is able to identify and classify relevant papers from a large and varied corpus Figure 5, starting from only a few examples. Our method has also shown that authors are often willing to provide useful clinical and contextual information about their work.

## CONCLUSION

This paper describes the development of a pipeline to retrieve papers containing COVID-19 immune signatures and for their semi-automated curation. Within this pipeline, we incorporated machine learning to classify papers from a bootstrapped sample of an existing repository (CORD-19). We found that SPECTER embedding provides a good reduced representation of a paper and its relatedness to other papers that can be adopted for the purpose of identifying scientifically salient features of the paper (in this case immune signatures). However, SPECTER was not a necessary component, as TF-IDF with logistic regression has similar performance to the SPECTER approach. Thirty-one percent of authors of papers with immune signatures voluntarily provided semistructured representations in response to a request from our team, regardless of the immune signature type.

Given its start as a neuroinformatics tool, the successful application to COVID-19 demonstrates that this pipeline approach is readily adaptable for other fields to identify papers containing scientifically relevant features, which can be further processed – by data solicitation, manual curation, or automated means – to extract the relevant data for presentation in a unified knowledge base or dashboard.

## ACKNOWLEDGMENTS

We thank Daniel Chawla and Bram Gerritsen for reviewing papers and providing feedback about the pipeline interface.

## SUPPLEMENTARY MATERIAL

**Table 1.** Abstracts containing any of the terms listed would be given the corresponding tag. Reviews can see these tags and search or filter abstracts with them.

| Tag | Terms |
| --- | --- |
| brain | brain, neurological, neuron, neurons |
| heart | heart, cardiac |
| kindey | kidney |
| liver | liver |
| lung | lung, pulmonary |
| artery | artery |
| skin | skin |
| bone | bone |
| upregulated | upregulated, up-regulated |
| downregulated | downregulated, down-regulated |
| rnaseq | rnaseq, rna-seq |
| immune response | immune response |
| t-cell | t cell, t cells, t-cell, t-cells |
| b-cell | b cell, b cells, b-cell, b-cells |
| virus | virus, viral |
| human | human, humans, patient, patients |
| antibody | antibody, antibodies |
| treatment | treatment, medicine |
| mouse | mouse, mice |
| macrophage | macrophage, macrophages |
| protein | protein, proteins |
| gene | gene, genes, genetic |

**Table 2.**
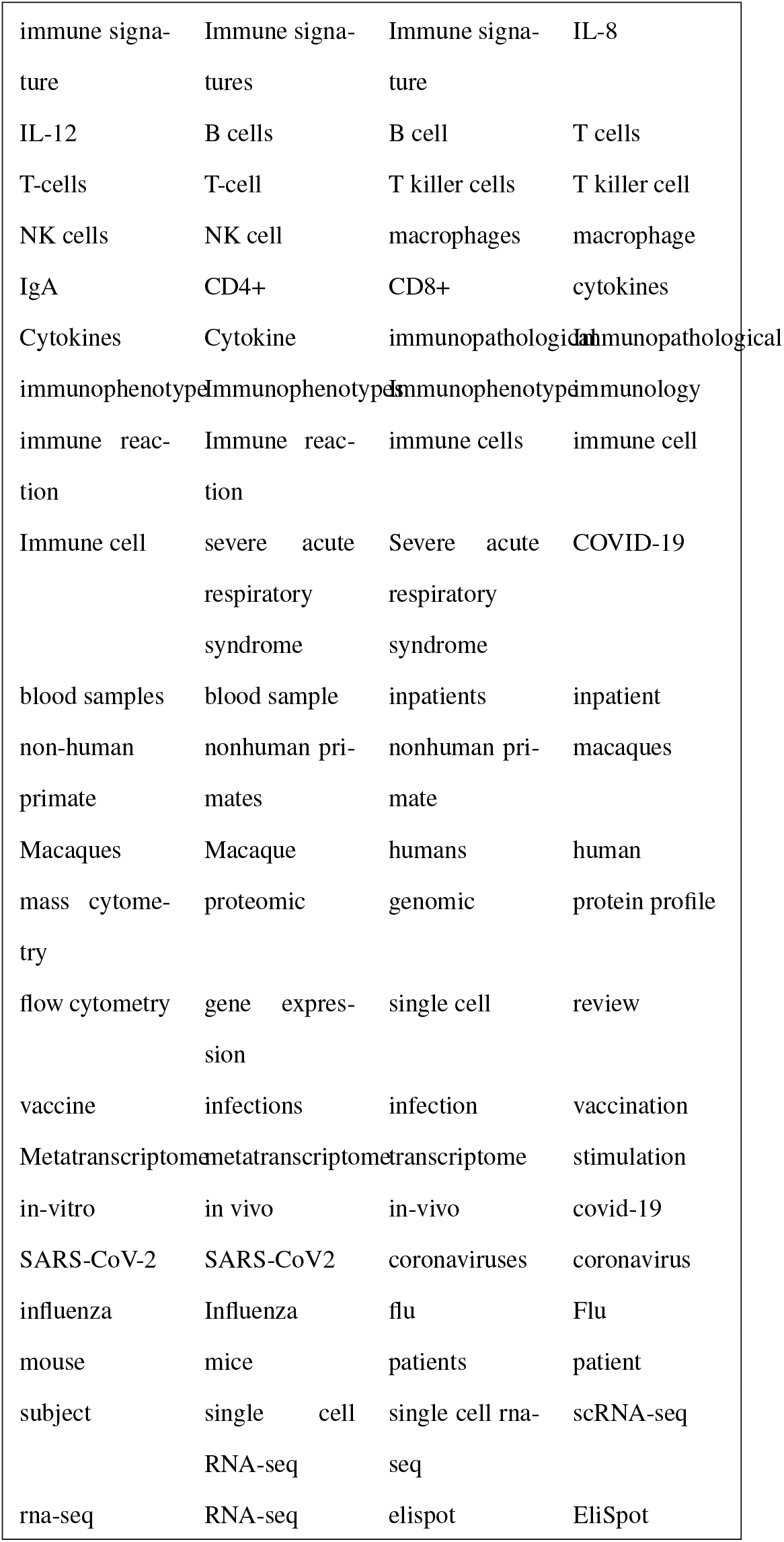
Terms identified by a reviewer that we highlight in the interface to facilitate classification.

**Table 3.** Differences in relative frequencies between papers predicted to have COVID-19 immune signatures and the signature type. The ten words with largest differences in relative word frequencies are shown. Bold text indicates the word had a higher relative frequent in the set, light text that it was less frequent. Relative frequencies are given as the occurrences of the word as a percent of the corpus.

| words | relative frequency (%) |  |
| --- | --- | --- |
|  | with immune signatures | without immune signatures |
| **patient** | **6.45** | **0.98** |
| pandemic | 0.13 | 0.54 |
| **severe** | **1.92** | **0.16** |
| health | 0.12 | 0.29 |
| **cell** | **1.20** | **0.11** |
| **severity** | **0.86** | **0.05** |
| review | 0.01 | 0.13 |
| **blood** | **0.67** | **0.02** |
| **level** | **1.24** | **0.13** |
| testing | 0.04 | 0.16 |
|  | with type A signatures | with type B or C signatures |
| patient | 3.10 | 14.21 |
| clinical | 0.46 | 3.35 |
| hospital | 0.05 | 2.07 |
| covid | 5.24 | 11.10 |
| group | 0.19 | 2.39 |
| **gene** | **1.91** | **0.04** |
| admission | 0.03 | 1.72 |
| mortality | 0.17 | 2.21 |
| laboratory | 0.02 | 1.25 |
| risk | 0.20 | 1.81 |
|  | with type B signatures | with type A or C signatures |
| gene | 0.03 | 1.63 |
| covid | 4.65 | 9.31 |
| cell | 1.22 | 3.26 |
| patient | 5.66 | 9.13 |
| expression | 0.09 | 0.95 |
| lung | 0.11 | 0.94 |
| immune | 0.45 | 1.58 |
| pathway | 0.05 | 0.66 |
| injury | 0.25 | 1.08 |
| sars | 0.80 | 2.02 |
|  | with type C signatures | with type A or B signatures |
| cell | 0.09 | 4.34 |
| immune | 0.05 | 1.96 |
| response | 0.09 | 1.68 |
| sars | 0.39 | 2.40 |
| covid | 4.78 | 9.02 |
| severe | 0.90 | 2.89 |
| infection | 0.35 | 1.69 |
| cytokine | 0.03 | 0.85 |
| pathway | 0.01 | 0.70 |
| disease | 1.24 | 3.05 |

## REFERENCES

1. National Science Foundation National Science Board. Publication output: Us trends and international comparisons. Science and Engineering Indicators, 2019, 2020.

2. Ricardo Baeza-Yates, Berthier Ribeiro-Neto, et al. Modern information retrieval, volume 463. ACM press New York, 1999.

3. Shah Khalid, Shah Khusro, Irfan Ullah, and Godfrey Dawson-Amoah. On the current state of scholarly retrieval systems. Engineering, technology & applied science research, 9(1): 3863–3870, 2019.

4. Kirk Roberts, Tasmeer Alam, Steven Bedrick, Dina Demner-Fushman, Kyle Lo, Ian Soboroff, Ellen Voorhees, Lucy Lu Wang, and William R Hersh. Trec-covid: rationale and structure of an information retrieval shared task for covid-19. Journal of the American Medical Informatics Association, 27(9):1431–1436, 2020.

5. Kenneth C Smith, Daniel G Chawla, Bhavjinder K Dhillon, Zhou Ji, Randi Vita, Eva van der Leest, Jessica Weng, Ernest Tang, Amani Abid, Bjoern Peters, et al. A curated collection of human vaccination response signatures. bioRxiv, 2021.

6. T. M. Morse, R. Wang, N.T. Carnevale, G. M. Shepherd, and R. A. McDougal. Pipeline to promote discovery and sharing of computational neuroscience research. In Program Number 814.07 Neuroscience Meeting Planner. Society for Neuroscience, 2017.

7. Kevin L Howe, Premanand Achuthan, James Allen, Jamie Allen, Jorge Alvarez-Jarreta, M Ridwan Amode, Irina M Armean, Andrey G Azov, Ruth Bennett, Jyothish Bhai, et al. Ensembl 2021. Nucleic acids research, 49(D1):D884–D891, 2021.

8. UniProt Consortium. Uniprot: a worldwide hub of protein knowledge. Nucleic acids research, 47(D1):D506–D515, 2019.

9. Robert A McDougal, Isha Dalal, Thomas M Morse, and Gordon M Shepherd. Automated metadata suggestion during repository submission. Neuroinformatics, 17(3):361– 371, 2019.

10. Tian Kang, Shirui Zou, and Chunhua Weng. Pretraining to recognize pico elements from randomized controlled trial literature. Studies in health technology and informatics, 264:188, 2019.

11. Si Yang, Ying Wu, Yujiao Deng, Linghui Zhou, Pengtao Yang, Yi Zheng, Dai Zhang, Zhen Zhai, Na Li, Qian Hao, et al. Identification of a prognostic immune signature for cervical cancer to predict survival and response to immune checkpoint inhibitors. Oncoimmunology, 8(12):e1659094, 2019.

12. Lucy Lu Wang, Kyle Lo, Yoganand Chandrasekhar, Russell Reas, Jiangjiang Yang, Darrin Eide, Kathryn Funk, Rodney Kinney, Ziyang Liu, William Merrill, et al. Cord-19: The covid-19 open research dataset. ArXiv, 2020.

13. Ellen Voorhees, Tasmeer Alam, Steven Bedrick, Dina Demner-Fushman, William R Hersh, Kyle Lo, Kirk Roberts, Ian Soboroff, and Lucy Lu Wang. Trec-covid: constructing a pandemic information retrieval test collection. In ACM SIGIR Forum, volume 54, pages 1–12. ACM New York, NY, USA, 2021.

14. Jimmy S Chen and William R Hersh. A comparative analysis of system features used in the trec-covid information retrieval challenge. Journal of Biomedical Informatics, 117:103745, 2021.

15. Arman Cohan, Sergey Feldman, Iz Beltagy, Doug Downey, and Daniel S Weld. Specter: Document-level representation learning using citation-informed transformers. In Proceedings of the 58th Annual Meeting of the Association for Computational Linguistics, pages 2270–2282, 2020.

16. F. Pedregosa, G. Varoquaux, A. Gramfort, V. Michel, B. Thirion, O. Grisel, M. Blondel, P. Prettenhofer, R. Weiss, V. Dubourg, J. Vanderplas, A. Passos, D. Cournapeau, M. Brucher, M. Perrot, and E. Duchesnay. Scikit-learn: Machine learning in Python. Journal of Machine Learning Research, 12:2825–2830, 2011.

17. John Platt et al. Probabilistic outputs for support vector machines and comparisons to regularized likelihood methods. Advances in large margin classifiers, 10(3):61–74, 1999.

18. The pandas development team. pandas-dev/pandas: Pandas, February 2020.

19. Leland McInnes, John Healy, and James Melville. Umap: Uniform manifold approximation and projection for dimension reduction. arXiv preprint 1802.03426, 2018.

20. Pauli Virtanen, Ralf Gommers, Travis E Oliphant, Matt Haberland, Tyler Reddy, David Cournapeau, Evgeni Burovski, Pearu Peterson, Warren Weckesser, Jonathan Bright, et al. Scipy 1.0: fundamental algorithms for scientific computing in python. Nature methods, 17(3):261– 272, 2020.

21. Steven Bird. Nltk: the natural language toolkit. In Proceedings of the COLING/ACL 2006 Interactive Presentation Sessions, pages 69–72, 2006.

22. Christiane Fellbaum, editor. WordNet: An Electronic Lexical Database. Language, Speech, and Communication. MIT Press, Cambridge, MA, 1998. ISBN 978-0-262-06197-1.

23. Radim R?ehõ?rek and Petr Sojka. Software Framework for Topic Modelling with Large Corpora. In Proceedings of the LREC 2010 Workshop on New Challenges for NLP Frameworks, pages 45–50, Valletta, Malta, May 2010. ELRA. http://is.muni.cz/publication/884893/en.

24. David M Blei, Andrew Y Ng, and Michael I Jordan. Latent dirichlet allocation. the Journal of machine Learning research, 3:993–1022, 2003.

25. H. Akaike. A new look at the statistical model identification. Automatic Control, IEEE Transactions on, 19(6):716–723, 1974.

26. Jacob Cohen. A coefficient of agreement for nominal scales. Educational and psychological measurement, 20(1):37–46, 1960.

27. Karen Sparck Jones. A statistical interpretation of term specificity and its application in retrieval. Journal of documentation, 1972.

28. Paul Rayson and Roger Garside. Comparing corpora using frequency profiling. In The workshop on comparing corpora, pages 1–6, 2000.

29. Robert A McDougal, Thomas M Morse, Ted Carnevale, Luis Marenco, Rixin Wang, Michele Migliore, Perry L Miller, Gordon M Shepherd, and Michael L Hines. Twenty years of ModelDB and beyond: building essential modeling tools for the future of neuroscience. Journal of computational neuroscience, 42(1):1–10, 2017.

30. Giorgio A. Ascoli. Turning the tide of data sharing. Neuroinformatics, 17(4):473–474, 2019. doi: 10.1007/s12021-019-09437-8.

31. Waleed Ammar, Dirk Groeneveld, Chandra Bhagavatula, Iz Beltagy, Miles Crawford, Doug Downey, Jason Dunkelberger, Ahmed Elgohary, Sergey Feldman, Vu Ha, et al. Construction of the literature graph in semantic scholar. arXiv preprint 1805.02262, 2018.

32. James A Overton, Randi Vita, Patrick Dunn, Julie G Burel, Syed Ahmad Chan Bukhari, Kei-Hoi Cheung, Steven H Kleinstein, Alexander D Diehl, and Bjoern Peters. Reporting and connecting cell type names and gating definitions through ontologies. BMC bioinformatics, 20(5):259–264, 2019.

33. Naveen Tirupattur. Text miner for hypergraphs using output space sampling. Purdue University, 2011.

34. Benjamin Nye, Junyi Jessy Li, Roma Patel, Yinfei Yang, Iain J Marshall, Ani Nenkova, and Byron C Wallace. A corpus with multi-level annotations of patients, interventions and outcomes to support language processing for medical literature. In Proceedings of the conference. Association for Computational Linguistics. Meeting, volume 2018, page 197. NIH Public Access, 2018.

35. Vladimir Brusic, Raphael Gottardo, Steven H Kleinstein, and Mark M Davis. Computational resources for high-dimensional immune analysis from the human immunology project consortium. Nature biotechnology, 32(2):146–148, 2014.

36. Jernej Godec, Yan Tan, Arthur Liberzon, Pablo Tamayo, Sanchita Bhattacharya, Atul J Butte, Jill P Mesirov, and W Nicholas Haining. Compendium of immune signatures identifies conserved and species-specific biology in response to inflammation. Immunity, 44(1):194–206, 2016.

